# Systematic discovery of uncharacterized transcription factors in *Escherichia coli* K-12 MG1655

**DOI:** 10.1101/343913

**Authors:** Ye Gao, James T. Yurkovich, Sang Woo Seo, Ilyas Kabimoldayev, Andreas Dräger, Ke Chen, Anand V. Sastry, Xin Fang, Nathan Mih, Laurence Yang, Johannes Eichner, Byung-Kwan Cho, Donghyuk Kim, Bernhard O. Palsson

## Abstract

Transcriptional regulation enables cells to respond to environmental changes. Yet, among the estimated 304 candidate transcription factors (TFs) in *Escherichia coli* K-12 MG1655, 185 have been experimentally identified and only a few tens of them have been fully characterized by ChIP methods. Understanding the remaining TFs is key to improving our knowledge of the *E. coli* transcriptional regulatory network (TRN). Here, we developed an integrated workflow for the computational prediction and comprehensive experimental validation of TFs using a suite of genome-wide experiments. We applied this workflow to: 1) identify 16 candidate TFs from over a hundred candidate uncharacterized genes; 2) capture a total of 255 DNA binding peaks for 10 candidate TFs resulting in six high-confidence binding motifs; 3) reconstruct the regulons of these 10 TFs by determining gene expression changes upon deletion of each TF; and 4) determine the regulatory roles of three TFs (YiaJ, YdcI, and YeiE) as regulators of L-ascorbate utilization, proton transfer and acetate metabolism, and iron homeostasis under iron limited condition, respectively. Together, these results demonstrate how this workflow can be used to discover, characterize, and elucidate regulatory functions of uncharacterized TFs in parallel.

## INTRODUCTION

Transcription factors (TFs) modulate gene expression in response to environmental perturbations by interacting with a combination of sigma factors, RNA polymerase (RNAP), activating metabolites, and inorganic compounds. These signals collectively lead TFs to bind to specific DNA sequences referred to as binding motifs (1). Microorganisms therefore have the ability to quickly adapt to diverse and extreme environmental conditions. In the transcription regulation, genes are indirectly or directly regulated by one or more TFs. A set of genes directly controlled by the same TF are considered to belong to a regulon (2), with the complete set of regulons forming the transcriptional regulatory network (TRN).

Databases such as EcoCyc (3, 4), RegulonDB (5), and TEC (6) maintain large amounts of information about TFs. However, a complete TRN for individual organisms still does not exist due to challenges associated with: 1) the identification of all TFs; 2) characterizing transcription factor binding sites (TFBS); and 3) TRN reconstruction.

First, genome-scale annotation of genes is required for the identification of the complete set of TFs. The emergence of high-throughput DNA sequencing has created a large number of candidate protein-encoding DNA sequences, leading to an increased demand for the discovery and annotation of protein functions. However, assigning a physiological function to the sequenced yet uncharacterized genes is still a substantial challenge (7, 8). For example, although *Escherichia coli* K-12 MG1655 has one of the most widely-studied genomes, functional annotation is still missing for approximately 30% of its genes (9). This lack of annotation includes an estimated 50-80 uncharacterized TFs in *E. coli* K-12 MG1655 (6). The percentage of uncharacterized genes in other strains may even higher than that of a model prokaryote *E. coli* K-12. Thus, a new computational approach is needed to predict a complete set of TFs in prokaryotes.

Second, a genome-wide characterization of TFBS is essential for the reconstruction of a global TRN. Despite a large amount of knowledge about microbial TFs in databases and the literature, a lot of individual TFs remain to be studied about their binding sites. Traditionally, TFBS are identified through approaches such as DNase I footprinting and electromobility shift assays, which are limited to the interactions between TFs and single targets (10). With advances in genome-wide research technologies, many TFs have been experimentally investigated using the systematic evolution of ligands by exponential enrichment (SELEX) and chromatin immunoprecipitation with microarray (ChIP-chip) or by sequencing (ChIP-seq) (6, 11–16). Recently, the ChIP-seq protocol has been combined with an exonuclease treatment (ChIP-exo) to reflects *in vivo* regulatory interactions between TFs and target genes at a single-base-pair resolution (17). In addition, ChIP-exo can be easily applied to investigate differential binding patterns of the same TF under different environmental conditions (18–20). Thus, ChIP-exo provides us with a powerful approach to characterize TFBS at the genome-scale.

Third, several computational approaches have been developed for the reconstruction of the TRN, including the use of gene expression data (21, 22), regulon-based associations (23), and integrated analysis with metabolic models (24). The expression data-driven approach for TRN reconstruction was widely used to predict transcription factor activities in *E. coli* K-12. Recently, we have supplemented ChIP-exo with transcription profiling to describe the regulons of major TFs, including Cra, ArgR, Fur, OxyR, SoxRS, OmpR, and GadEWX (20, 25–29). Therefore, this well-described approach is successfully applied to TRN reconstruction.

Here, we address these three challenges through the development of an integrated computational and experimental workflow to discover uncharacterized TFs in prokaryotes. Using *E. coli* K-12 MG1655 as an example, we used the algorithm, TFpredict (30), that uses machine learning to computationally predict whether a given protein is a candidate TF. Given the resulting list of candidate TFs, we then examined their DNA-binding domains, predicted their active conditions, and performed an *in vivo* experimental validation of predicted DNA-binding capabilities. This workflow resulted in the elucidation of the biological functions of three uncharacterized TFs (YiaJ, YdcI, and YeiE) through an in-depth analysis of mutant phenotypes. Together, these results demonstrate the utility of our systematic identification workflow and provide a roadmap for its use in other organisms.

## MATERIAL AND METHODS

### Computational identification of candidate TFs

We modified the existing TFpredict algorithm that was designed for eukaryotic organisms to enable its use on prokaryotes. In brief, this algorithm takes a protein sequence as input and outputs a quantified score in the range [0,1] that represents the likelihood that it is a TF based on homology. A score of zero denotes that a protein is unlikely to be a TF; a score of one denotes that a protein is very likely to be a TF. Our modifications adapted the algorithm for use with prokaryotic protein sequences, instead of eukaryotic protein sequences. Before running the algorithm, we filtered the input protein sequences to only include those that were annotated as DNA-binding and “reviewed” in UniProt (31). We then filtered by GO terms to ensure that no node had a parent node with a GO term related to another class. Finally, we filtered the input proteins to exclude any protein sequences that were annotated with non-TF keywords: kinase, ubiquitin, actin, antigen, biotin, histone, chaperone, tubulin, transmembrane protein, endonuclease, exonuclease, translation initiation factor.

### Bacterial strains, media, and growth conditions

All strains used in this study are *E. coli* K-12 MG1655 and its derivatives, deletion strains and myc-tagged strains (Dataset S2). For ChIP-exo experiments, the *E. coli* strains harboring 8-myc were generated by a λ red-mediated site-specific recombination system targeting C-terminal region as described previously (32). For expression profiling by RNA-seq, deletion strains *ΔydcI, ΔyeiE, ΔyafC, ΔyiaJ, ΔyheO, ΔybaO, ΔybaQ, ΔybiH, ΔyddM* and *ΔyieP* were also constructed by a λ red-mediated site-specific recombination system (33). For ChIP-exo experiments, glycerol stocks of *E. coli* strains were inoculated into M9 minimal media (47.8 mM Na_2_H PO_4_, 22mM KH_2_PO_4_, 8.6 mM NaCl, 18.7 mM NH_4_Cl, 2 mM MgSO_4_ and 0.1 mM CaCl_2_) with 0.2% (w/v) glucose. M9 minimal media was also supplemented with 1 mL trace element solution (100X) containing 1 g EDTA, 29 mg ZnSO_4_.7H_2_O, 198 mg MnCl_2_.4H_2_O, 254 mg CoCl_2_.6H_2_O, 13.4 mg CuCl_2_, and 147 mg CaCl_2_. The culture was incubated at 37 °C overnight with agitation, and then was used to inoculate the fresh media (1/200 dilution). The volume of the fresh media was 150 mL for each biological replicate. The fresh culture was incubated at 37 °C with agitation to the mid-log phase (OD_600_ ≈ 0.5). For RNA-seq expression profiling, glycerol stocks of *E. coli* strains were inoculated into M9 minimal media with the same carbon sources as used in the ChIP-exo experiment for each TF candidate. The concentration of carbon sources was 0.2% (w/v). M9 minimal media was also supplemented with 1 mL trace element solution (100X). The culture was incubated at 37 °C overnight with agitation, and then was used to inoculate the fresh media. The fresh culture was incubated at 37 °C with agitation to the mid-log phase (OD_600_ ≈ 0.5).

### Measurement of bacterial growth

The effects of iron limited conditions on cell growth were examined by growing *E. coli* K-12 MG1655 and *yeiE* deletion strain under four media treatments: (1) M9 minimal glucose medium; (2) M9 minimal glucose medium containing 0.2 mM of the iron chelating agent 2,2’-dipyridyl (DPD) (Fluka); (3) M9 minimal glucose medium containing 0.3 mM of DPD; (4) M9 minimal glucose medium containing 0.4 mM DPD. Cells grown overnight on M9 minimal glucose medium at 37 °C with agitation were inoculated into these four kinds of fresh media. Aliquots of overnight cell culture were diluted 1: 200 into four kinds of fresh media, then were incubated at 37 °C with agitation. Growth was measured and recorded by OD_600_ using Thermo BIOMATE 3S UV-visible spectrophotometer.

Similarly, to measure growth rate on low pH or acetate medium, the culture was incubated at low pH or acetate medium at 37 °C overnight with agitation, and then was used to inoculate the fresh media (1/200 dilution). The volume of the fresh media was 150 mL. The fresh culture was incubated at 37 °C with agitation. Growth was measured and recorded by OD_600_ using Thermo BIOMATE 3S UV-visible spectrophotometer.

To measure growth on L-ascorbate, cells were grown anaerobically in medium containing L-ascorbate as described by the literature (34). Briefly, *E. coli* strains were grown overnight on M9 minimal glucose medium, and the cells were suspended in M9 salts medium. The aliquots were adjusted to 1.0, and 100 ul aliquots were inoculated into 10 mL culture tubes (Fisher Scientific) that were filled to the top with M9 minimal medium with a concentration of 20 mM L-ascorbate. Then the culture tubes were capped, sealed with parafilm, and then incubated at 37 °C. Six independent experiments were conducted.

### ChIP-exo experiment

ChIP-exo experimentation was performed following the procedures previously described (35). In brief, to identify each TF candidate binding maps *in vivo*, we isolated the DNA bound to each TF candidate from formaldehyde cross-linked *E. coli* cells by chromatin immunoprecipitation (ChIP) with the specific antibodies that specifically recognize myc tag (9E10, Santa Cruz Biotechnology), and Dynabeads Pan Mouse IgG magnetic beads (Invitrogen) followed by stringent washings as described previously (36). ChIP materials (chromatin-beads) were used to perform on-bead enzymatic reactions of the ChIP-exo method (37). Briefly, the sheared DNA of chromatin-beads was repaired by the NEBNext End Repair Module (New England Biolabs) followed by the addition of a single dA overhang and ligation of the first adaptor (5’-phosphorylated) using dA-Tailing Module (New England Biolabs) and NEBNext Quick Ligation Module (New England Biolabs), respectively. Nick repair was performed by using PreCR Repair Mix (New England Biolabs). Lambda exonuclease- and RecJ_f_ exonuclease-treated chromatin was eluted from the beads and the protein-DNA cross-link was reversed by overnight incubation at 65°C. RNAs- and Proteins-removed DNA samples were used to perform primer extension and second adaptor ligation with following modifications. The DNA samples incubated for primer extension as described previously (35) were treated with dA-Tailing Module (New England Biolabs) and NEBNext Quick Ligation Module (New England Biolabs) for second adaptor ligation. The DNA sample purified by GeneRead Size Selection Kit (Qiagen) was enriched by polymerase chain reaction (PCR) using Phusion High-Fidelity DNA Polymerase (New England Biolabs). The amplified DNA samples were purified again by GeneRead Size Selection Kit (Qiagen) and quantified using Qubit dsDNA HS Assay Kit (Life Technologies). Quality of the DNA sample was checked by running Agilent High Sensitivity DNA Kit using Agilent 2100 Bioanalyzer (Agilent) before sequenced using HiSeq 2500 (Illumina) in accordance with the manufacturer’s instructions. Each modified step was also performed in accordance with the manufacturer’s instructions. ChIP-exo experiments were performed in biological duplicate.

### RNA-seq expression profiling

Three milliliters of cells from mid-log phase culture were mixed with 6 mL RNAprotect Bacteria Reagent (Qiagen). Samples were mixed immediately by vortexing for 5 seconds, incubated for 5 minutes at room temperature, and then centrifuged at 5000*g*, for 10 minutes. The supernatant was decanted and any residual supernatant was removed by inverting the tube once onto a paper towel. Total RNA samples were then isolated using RNeasy Plus Mini kit (Qiagen) in accordance with the manufacturer’s instruction. Samples were then quantified using a NanoDrop 1000 spectrophotometer (Thermo Scientific) and quality of the isolated RNA was checked by running RNA 6000 Pico Kit using Agilent 2100 Bioanalyzer (Agilent). Paired-end, strand-specific RNA-seq library was prepared using KAPA RNA Hyper Prep kit (KAPA Biosystems), following the instruction (38, 39). Resulting libraries were analyzed on an Agilent Bioanalyzer DNA 1000 chip (Agilent). Sequencing was performed on a Hiseq 2500 sequencer at the Genomics Core facility of University of California, San Diego.

### Peak calling for ChIP-exo dataset

Peak calling was performed as previously described (35). Sequence reads generated from ChIP-exo were mapped onto the reference genome (NC_000913.2) using bowtie (40) with default options to generate SAM output files (Dataset S3). MACE program (41) was used to define peak candidates from biological duplicates for each experimental condition with sequence depth normalization. To reduce false-positive peaks, peaks with signal-to-noise (S/N) ratio less than 1.5 were removed. The noise level was set to the top 5% of signals at genomic positions because top 5% makes a background level in plateau and top 5% intensities from each ChIP-exo replicates across conditions correlate well with the total number of reads (35, 42, 43). The calculation of S/N ratio resembles the way to calculate ChIP-chip peak intensity where IP signal was divided by Mock signal. Then, each peak was assigned to the nearest gene. Genome-scale data were visualized using MetaScope (http://systemsbiology.ucsd.edu/Downloads/MetaScope).

### Motif search from ChIP-exo peaks

The sequence motif analysis for TFs and σ-factors was performed using the MEME software suite (44). For YdcI, YbiH, YbaQ, YeiE, YddM and YieP, sequences in binding regions were extracted from the reference sequence (NC_000913.2).

### Calculation of differentially expressed gene

Sequence reads generated from RNA-seq were mapped onto the reference genome (NC_000913.2) using bowtie (40) with the maximum insert size of 1000 bp, and 2 maximum mismatches after trimming 3 bp at 3’ ends (Dataset S4). SAM files generated from bowtie were then used for Cufflinks (http://cufflinks.cbcb.umd.edu/) (45) to calculate fragments per kilobase of exon per million fragments (FPKM). Cufflinks was run with default options with the library type of dUTP RNA-seq and the default normalization method (classic-fpkm). Expression with log_2_ fold change ≥ log_2_(1.5) and *q*-value ≤ 0.05 or log_2_fold change≤ -log_2_(1.5) and *q*-value ≤ 0.05 was considered as differentially expressed. Genome-scale data were visualized using MetaScope.

### COG functional enrichment

The regulons were categorized according to their annotated clusters of orthologous groups (COG) category (46). Functional enrichment of COG categories in the target genes was determined by performing hypergeometric test, and *p-value* < 0.05 was considered significant.

### Structural analysis of candidate TFs

Homology models of the candidate transcription factors YdcI, YeiE, and YiaJ were constructed using the SWISS-MODEL pipeline, which also carries out a prediction of the oligomeric state of the enzyme (47). Multiple templates were analyzed and inference of the oligomeric state was based on the reported interface conservation scores to existing complexes of similar sequence identity. The structures were annotated using information in UniProt (31) and visualized with VMD (48).

### Phylogenetic tree analysis

We queried the NCBI database for the homologs sequences of candidate TF YdcI across typical gram-negative strains, in order to show the shared origin of them. The phylogenetic tree (Neighbour-joining without distance corrections) was generated by MUSCLE (49).

## RESULTS

### Establishing a workflow to discover uncharacterized transcription factors

Our goal was to define a workflow that would computationally identify and experimentally validate candidate TFs at the genome-scale. To prioritize a list of uncharacterized TFs for experimental validation, we first reviewed and filtered the proteins in the *E. coli* K-12 MG1655 genome that met the following criteria: (1) the protein sequences were reviewed as non-redundant and authentic; (2) the proteins were annotated by Gene Ontology (50); (3) the proteins were annotated as non-TF; and (4) the proteins have functional annotation for DNA-binding or non-DNA binding (Supplementary Figure S1 and Dataset S1). This initial screening resulted in a unique set of 942 genes (of 4,460 total *E. coli* K-12 MG1655 genes) that were candidate TFs (Figure 1A).

**Figure 1.**
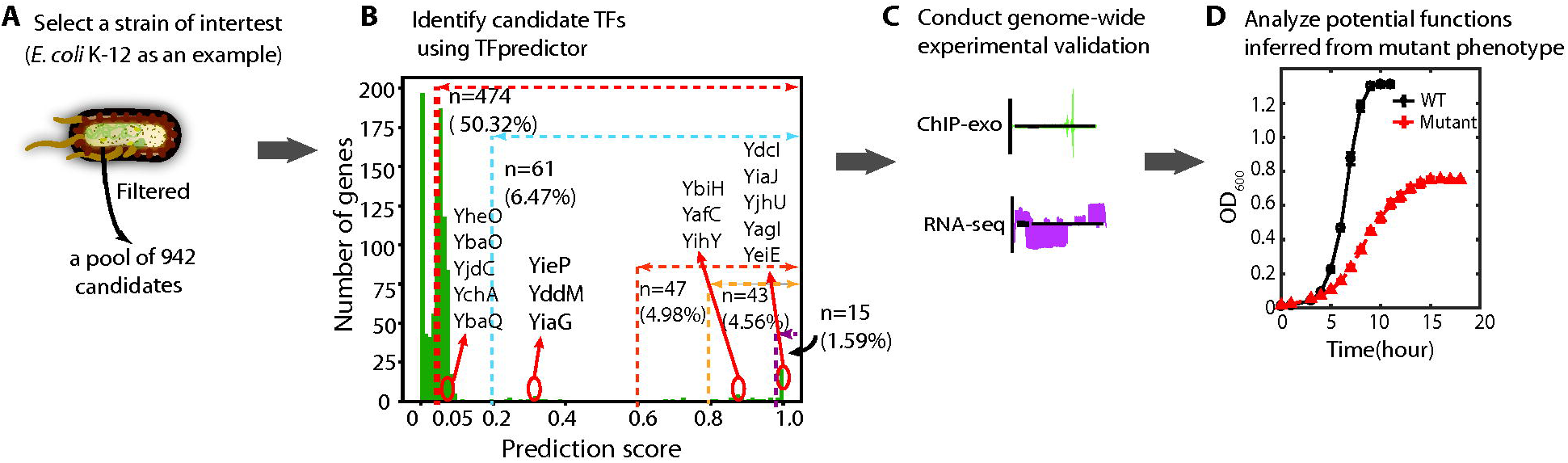
The scheme of the systematic workflow for identifying uncharacterized transcription factors in *E. coli* K-12 MG1655. (A) A general approach to identify candidate TFs in a bacterial strain of interest. Using *E. coli* K-12 MG1655 as an example, a pool of 942 uncharacterized proteins were identified after filtering. (B) Distribution of TFpredictor likelihood scores for the 942 uncharacterized candidate TF proteins (y-gene encoded). We empirically split prediction scores into 5 ranges separated by dashed lines: 1 (to the right of purple dashed line), [0.8,1) (yellow), [0.6,0.8) (orange), [0.2,0.6) (cyan), [0.05,0.2) (red). The number of candidate genes in each range and its percentage out of the total number of 942 uncharacterized proteins are indicated accordingly. The 16 candidate TFs selected for experimental validation are labeled within their score range. (C) Genome-wide experimental validation was conducted by identifying candidate TF binding to DNA sequence by ChIP-exo, and analyzing differential expression of their target genes by RNA-seq. (D) Hypothesized functions of selected candidate TFs were inferred by comparing phenotypes between WT and TF knockout mutants.

We then input these protein sequences into TFpredict (30), a machine learning algorithm that uses homology to predict whether a given eukaryotic protein is a TF. We modified this algorithm for use with prokaryotes (see Methods) and used it to rank the uncharacterized candidates in *E. coli* K-12 MG1655 based on their likelihood of being a TF (Figure 1B). Approximately 50% of the total uncharacterized candidates had prediction scores higher than 0.05 (Supplementary Table S1 and Figure S2). To provide insight into how well such a prediction score can guide discovery of new TFs, we considered candidates with scores in four different ranges: [0.05, 0.2), [0.2, 0.6), [0.8, 1), [1] for further experimental validation. 16 candidates came from these ranges: 1) range [1]: YdcI, YiaJ, YjhU, YagI, YeiE; 2) range [0.8, 1): YbiH, YafC, YihY; 3) range [0.2, 0.6): YieP, YddM, YiaG; 4) range [0.05, 0.2): YheO, YbaO, YjdC, YchA and YbaQ.

Next, to examine if selected candidates have DNA-binding peaks at the genome-scale, we conducted ChIP-exo experiments for each candidate (Figure 1C). Because uncharacterized TFs are likely to be expressed at low levels, especially at non-active conditions (51), it is necessary to determine the conditions under which uncharacterized TFs are active. We can infer the activating conditions for TFs based on their biochemical functions, since TFs are activated by specific molecules or environments. We are thus able to classify 16 candidates into three groups based on the level of confidence of their biochemical/biological roles (Table 1). The first group contains candidates whose biochemical activities were studied *in vitro* with gel shift assays (52) or SELEX (6, 53), yet their *in vivo* biological functions remain largely unknown. The second group consists of candidates whose biological functions could be predicted according to well-studied functions of homologous genes in a closely related strain. The third group includes candidates with neither biochemical characterization nor biological function prediction.

We then applied different approaches to predict active conditions for each group of candidate TFs through ChIP-exo experiments. For the first group, we inferred conditions based on biochemical features, e.g., a previous study showed that *yiaJ* might be involved in the catabolism of rare carbon sources (54). The conditions for the second group were inferred from functional studies in a closely related strain. For example, *ydcI* is a highly conserved gene and is responsible for pH stress response in *Salmonella enterica* serovar Typhimurium (55); thus it is likely to function at similar conditions in *E. coli* K-12, though it might play multiple biological roles in *E. coli*. Finally, conditions were inferred for the third group based on expression profile data from the NCBI GEO repository (56, 57) (Supplementary Figure S3). If the expression level of the candidate TF is relatively high under a certain condition, it might be inferred as a test condition. The data showed that yeiE is highly expressed in glucose medium compared to other conditions. In this study, we analyzed the characteristics of each candidate and predicted active conditions for the ChIP-exo experiments.

Next, for those candidates having DNA-binding sites, we further investigated expression profiles upon deletion of each TF. Combing DNA bindings from ChIP-exo with gene expression, we can form hypotheses for the regulatory functions of candidate TFs. Lastly, to test the hypotheses, we explored mutant phenotypes under active conditions (Figure 1D).

### Capturing a genome-wide distribution of uncharacterized transcription factors (TFs)

To validate the *in silico* predictions of candidate TFs, we employed ChIP-exo to determine the *in vivo* genome-wide DNA-binding events of each candidate during growth at active conditions. We then examined global binding profiles for all candidates using the peak calling algorithm MACE (58) and confirmed that 10 out of 16 were DNA-binding proteins (Figure 2A). A total of 255 reproducible binding peaks were identified for 241 unique binding sites (Dataset S5). Of the six unconfirmed candidates, YagI and YjhU had high prediction scores (score >0.8). Therefore, it is possible that these proteins are TFs, but are not active under the basal condition we used here.

**Figure 2.**
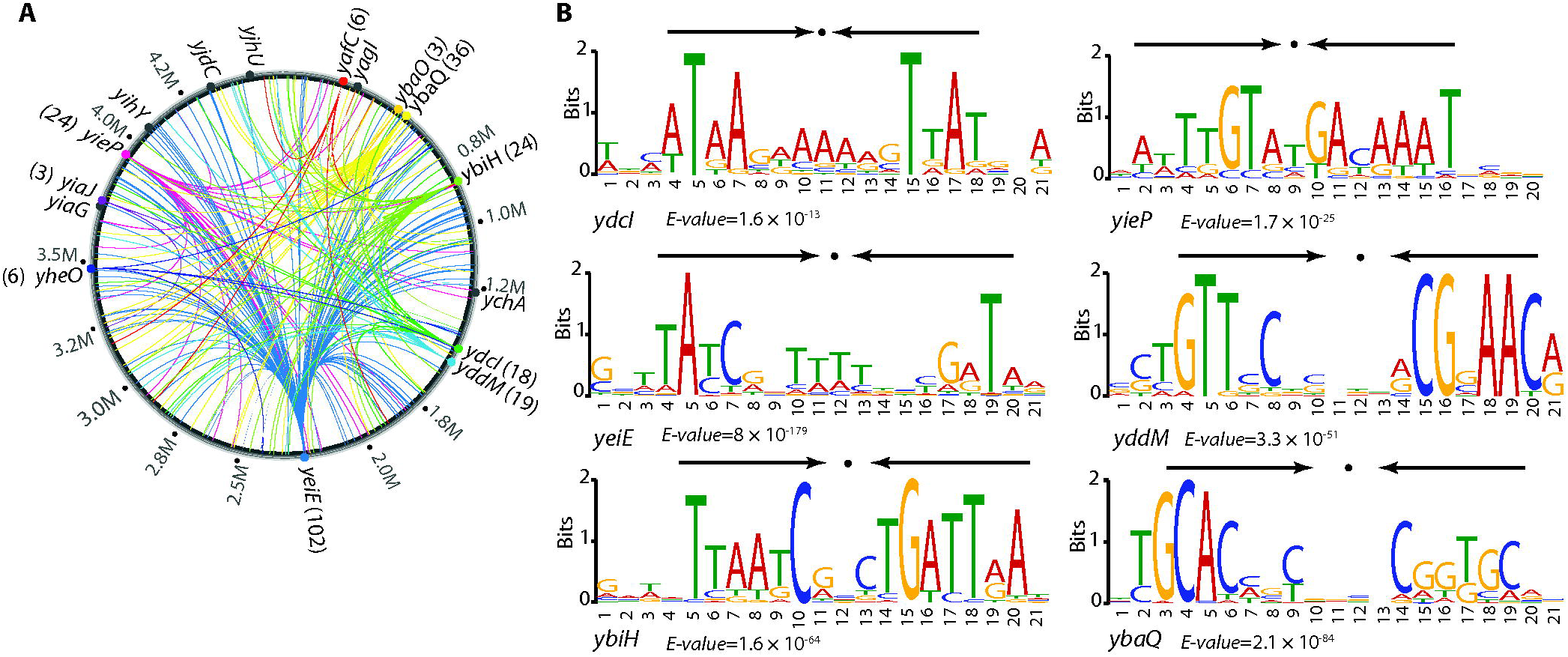
A global landscape of DNA binding events for uncharacterized TFs during growth at active conditions. (A) Binding sites identified by ChIP-exo. Verified uncharacterized TFs were labeled with colored circles. The numbers in the parentheses represent the number of identified binding sites for individual uncharacterized TFs. Gray circles represent uncharacterized TFs without binding peaks under the growth conditions used, which include YjhU, YjdC, YihY, YiaG, YagI, and YchA. (B) The sequence motifs for six uncharacterized TFs. The height of the letters (in bits on the y-axis) represents the degree of conservation at a given position within the aligned sequence set, with perfect conservation being 2 bits. Arrows above motif indicate the presence of palindromic sequence.

Compared to known global TFs, these ten uncharacterized TFs exhibit some interesting regulatory features. First, they have more intragenic binding peaks and fewer peaks located within putative regulatory regions. The binding sites from these confirmed TFs showed that only 41% (98 of 241) were located in putative regulatory regions (promoters and 5’-proximal to coding regions). Second, individual uncharacterized TFs had fewer binding peaks than those of global TFs such as CRP, Lrp, Fnr, and ArcA (12, 59, 60). Most of the uncharacterized TFs have 3-25 binding sites under active conditions, while global TFs in *E. coli* usually have more than 40 binding sites. Third, the uncharacterized TFs bind to more genes with putative functions (Supplementary Figure S4). Finally, the average expression level of these uncharacterized TFs are relatively lower than global TFs. These observations are consistent with the previous study showing that TF position in the TRN hierarchy network is correlated with its expression levels, its number of target genes, and its scope of regulatory function (61). TFs in the top hierarchy usually have high protein concentration in the cell, and regulate a large amount of genes of diverse functions. On the contrary, these candidate TFs are likely located in the lower levels of the *E. coli* hierarchical TRN, and may regulate local specific physiological functions instead of broad biological roles.

Next, using the MEME algorithm (62), we found conserved binding motifs for six of the 10 confirmed TFs (E-value < 10^−10^) (Figure 2B). Interestingly, the consensus binding motifs were palindromic, suggesting a dimeric protein conformation. Specifically, the transcriptional factor binding sites (TFBS) of YdcI and YbiH enclose AT-rich inverted repeats separated by 7-nt. This finding is consistent with the structural predictions (Supplementary Table S2 and Figure S5) that these TF candidates likely form homodimers or tetramers, which facilitate tight binding to DNA molecules in the cell.

### Interactions between uncharacterized TFs and RNA polymerase (RNAP)

A transcriptional repressor down-regulates transcription by steric exclusion of RNAP from the promoter regions. To determine the interaction between the uncharacterized TFs and RNAP, we compared the binding sites of the uncharacterized TFs with the −10 and −35 promoter elements occupied by RNAP. Three interactions modes are observed based on the relative location: 1) downstream (D) mode where TF binds downstream of the −10 and −35 promoter region (Figure 3A); 2) upstream (U) mode where TF binds upstream of the −10 and −35 promoter region (Figure 3B); and 3) overlap mode (O) where TF binding site coincides with the −10 and −35 promoter region (Figure 3C). To further illustrate how different TF-RNAP interaction modes may affect TF function, we characterize the regulatory effects on the target genes by their differential expression in ΔTF strain with respect to WT.

**Figure 3.**
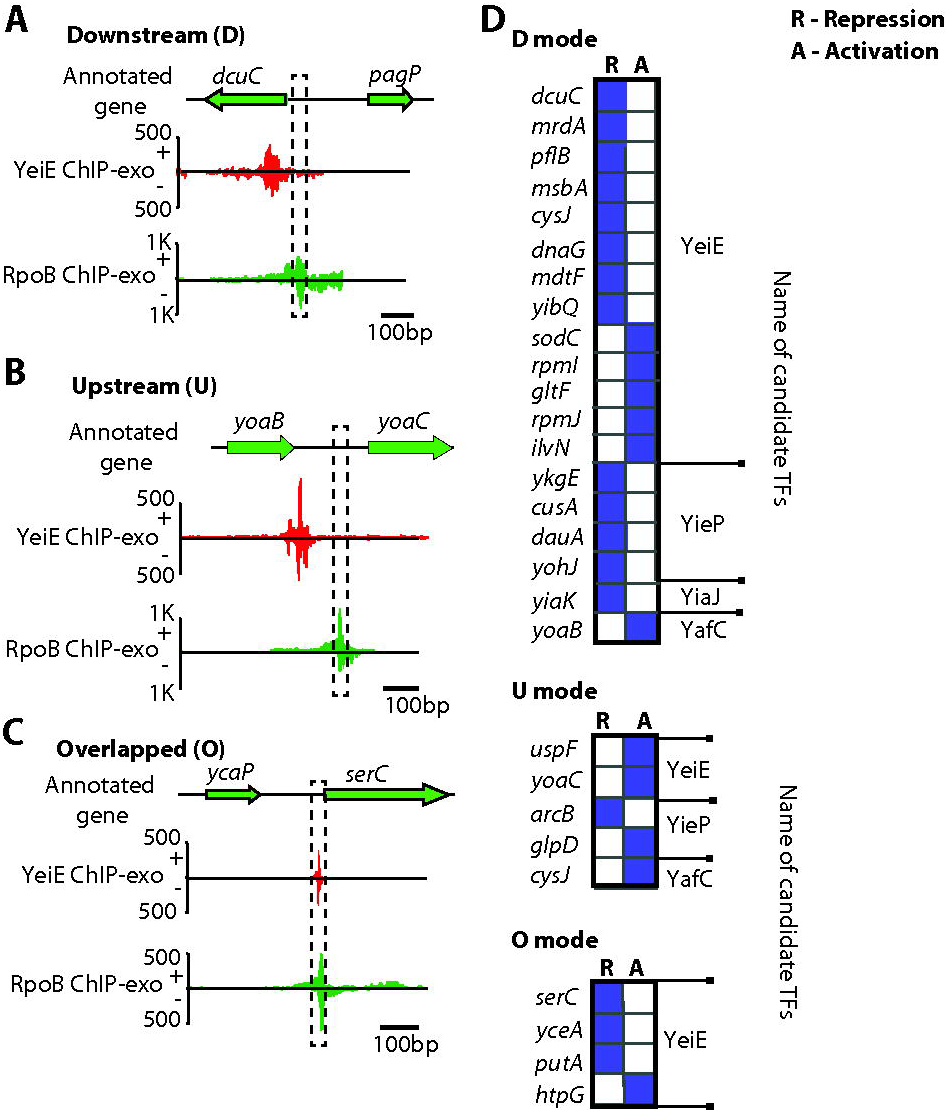
Transcriptional regulation by the position of uncharacterized TFs relative to the binding of RNA polymerase (RNAP), using binding sites from YeiE, YieP, YiaJ, and YafC as representatives. (A) In the case of *dcuC*, YeiE-binding is located at the downstream region of the promoter. (B) YeiE binds to the upstream site of the *yoaC*. (C) YeiE-binding region at the upstream of *serC* overlaps with the promoter occupied by RNAP. (D) The binding positions of YeiE, YieP, YiaJ, and YafC from the promoter are categorized according to the gene regulation. The abbreviations, D, U, and O indicate the downstream, upstream, and overlapped position, respectively. R, A indicate the regulation modes: repression and activation, respectively.

To demonstrate how binding sites of uncharacterized TFs interact with RNAP *in vivo*, we used four candidate TFs (YeiE, YieP, YiaJ, YafC) as representatives, since they showed a large number of binding sites. The most common binding mode for these transcription factors is downstream of the RNAP binding region. This binding mode commonly results in the repression of the target gene (13/19 or 68%). For example, YeiE binds downstream of the RNAP binding region of the gene *dcuC* and represses this gene (Figure 3A). However, the upstream binding mode is more commonly activation, as shown by *yoaC* (Figure 3B). Three of the four binding sites that overlap with RNAP binding location lead to the repression of the target genes *serC*, *yceA*, and *putA* (Figure 3C). Such genes as having the upstream, downstream or overlap modes from these four aforementioned representatives were determined (Figure 3D). This data suggested that transcriptional regulation by uncharacterized TFs are likely mediated by using steric exclusion mechanisms, though this pattern is not always true, as in *gltF*, *rpmI*, *ilvN*, and *htpG*. It is possible that other TFs are involved in the regulation of these target genes (Supplementary Figure S6) (63–66). Together, these data demonstrated that different sets of uncharacterized TFs have similar regulatory mechanisms, though they may have different biological functions.

To confirm the regulatory roles of candidate TFs, three of ten candidates identified by ChIP-exo (YiaJ, YdcI, YeiE) from three different groups were selected for further analysis, respectively. These three case studies illustrate how experimental observations from ChIP-exo and RNA-seq can be used to infer regulatory functions of a candidate TF. The binding sites of YiaJ and YdcI directly indicated the potential functions, so we used mutant phenotypes to validate biological roles. The genome-wide binding sites for YeiE showed that it is involved in diverse biological processes. Therefore we further combined expression profiling data with ChIP-exo to infer its potential roles in addition to mutant phenotype validation.

### Case I: YiaJ regulates genes that are responsible for the utilization of L-ascorbate

Group I contains candidates whose biochemical activities were studied *in vitro*, yet their *in vivo* biological functions still remain unclear. One of candidates is YiaJ, which has been studied *in vitro* by gel mobility shift assays (34, 67). However, *in vivo* analysis of direct interactions between YiaJ and DNA in *E. coli* has not been reported. In this study, we found that there were two binding peaks between the *yiaJ* and *yiaKLMNOPQRS* (*yiaK-yiaL-yiaM-yiaN-yiaO-lysK-sgbH-sgbU-sgbE*) operon (Figure 4A and Supplementary Figure S7). One binding peak suggested autogenous regulation and the other showed that YiaJ binds to a promoter region of the *yiaK-S* operon, which occupied the position of RNAP. We compared the expression data of the *yiaK-S* operon in the wild type and *yiaJ* deletion strain (Figure 4B) and found that the expression of the operon *yiaK-S* was highly up-regulated in the deletion strain. This result suggests the repression function of YiaJ on the *yiaK-S* operon. A previous study showed that YiaJ might be involved in the utilization of an uncommon carbon sugar (54). To further identify the substrate the *yiaK-S* operon catablizes, we compared the products of the *yiaK-S* operon with the known operon *ulaABCDEF* encoding for catabolic enzymes in the utilization of L-ascorbate, and found that the *yiaK-S* operon encodes similar catabolic enzymes in the L-ascorbate degradation pathway. Thus we proposed the regulatory role of YiaJ in *E. coli*, based on the products of the *yiaK-S* operon (Figure 4C). When L-ascorbate is imported and converted to L-ascorbate-6-phosphate by the PTS system in *E. coli* K-12 MG1655, expression of YiaJ would be repressed. Subsequently, *lyxK*, *sbgH*, *sgbU*, and *sgbE* encode four metabolic enzymes, L-xylulose kinase, gulonate-6- phosphate, L-xyluose-5-phosphate-3-epime, and L-ribulose-5-phosphate-4-epime, respectively. They can subsequently metabolize L-ascorbate-6-phosphate to D-xylulose-5-phosphate. Thus, *E. coli* could ferment L-ascorbate using a branch of the pentose metabolic pathway (34).

**Figure 4.**
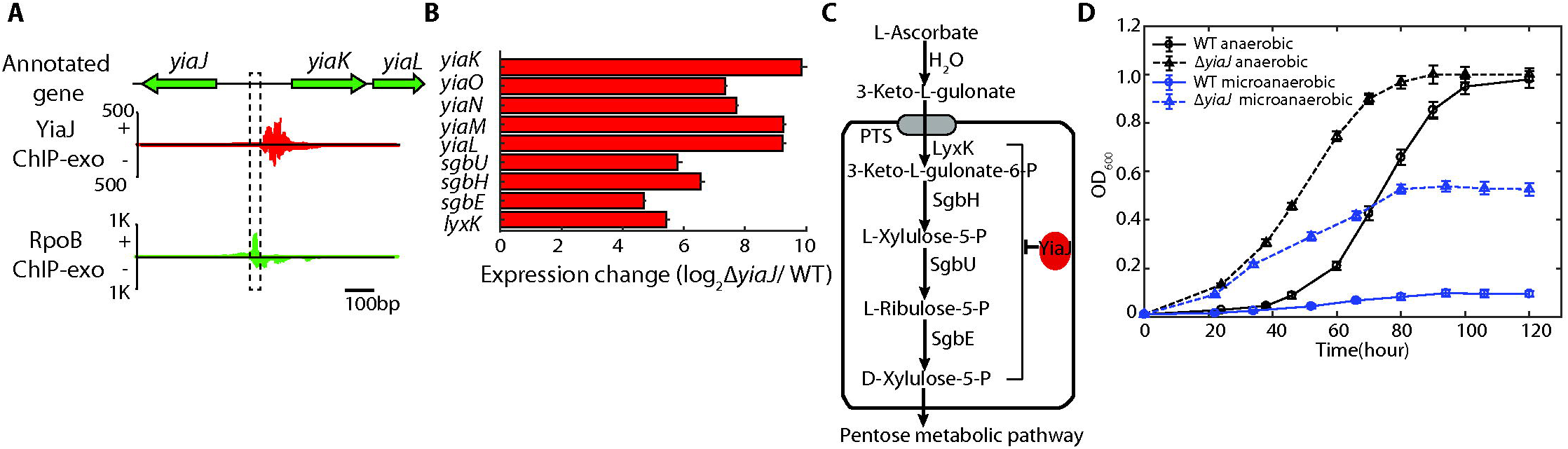
The regulatory role of the uncharacterized TF YiaJ is involved in utilization of L-ascorbate in *E. coli* K-12 MG1655. (A) YiaJ binding sites at the promoter region between *yiaJ* and the *yiaKLMNO-lyxK-sgbH-sgbU-sgbE* operon. (B) Expression changes for genes in the *yiaJ* deletion strain in the *yiaKLMNO-lyxK-sgbH-sgbU-sgbE* operon compared to the wild type strain. (C) The proposed function of YiaJ is to repress the ascorbate utilization pathway, therefore regulating the level of D-xylulose-5-P that feeds into the pentose phosphate pathway. (D) Growth curve of wild type and *yiaJ* deletion strains at ascorbate as the carbon source under anaerobic and microaerobic conditions, respectively.

To verify the function of the repressor YiaJ, we measured the growth profiles of the wild type and the *yiaJ* deletion strain in L-ascorbate medium. We found that the deletion of gene *yiaJ* allowed more rapid utilization of L-ascorbate and reduced the lag phase compared to wild type (Figure 4D). Furthermore, we found that the *yiaJ* deletion strain allowed cells to grow on L-ascorbate medium under microaerobic conditions, while the wild type could not. This confirmed that YiaJ is a repressor of operon *yiaK-S* and that it influenced growth under microaerobic conditions.

### Case II: YdcI is a transcription factor involved in proton and acetate metabolism

Group II consists of highly conserved candidate TFs, which were studied in a close related species. The regulatory function of the LysR-type regulator YdcI in *E. coli* K-12 MG1655 has not been studied with experimental approaches (6). Thus, we constructed a *ydcI* myc-tagged strain, and detected 18 binding sites using ChIP-exo (Supplementary Figure S8).

Previous studies showed that YdcI is responsible for acid stress and oxidative stress in *Salmonella enterica (55).* We analyzed the protein identity of YdcI among multiple strains across Gram-negative bacteria and found that YdcI is a highly and broadly conserved protein (Figure 5A). Notably, YdcI from *E. coli* K-12 MG1655 shares 80% of its identity with that from *Salmonella enterica*. Given that the function of a protein is tightly associated with its sequence, we can hypothesize that YdcI has similar biological roles in *E. coli* K-12 MG1655.

**Figure 5.**
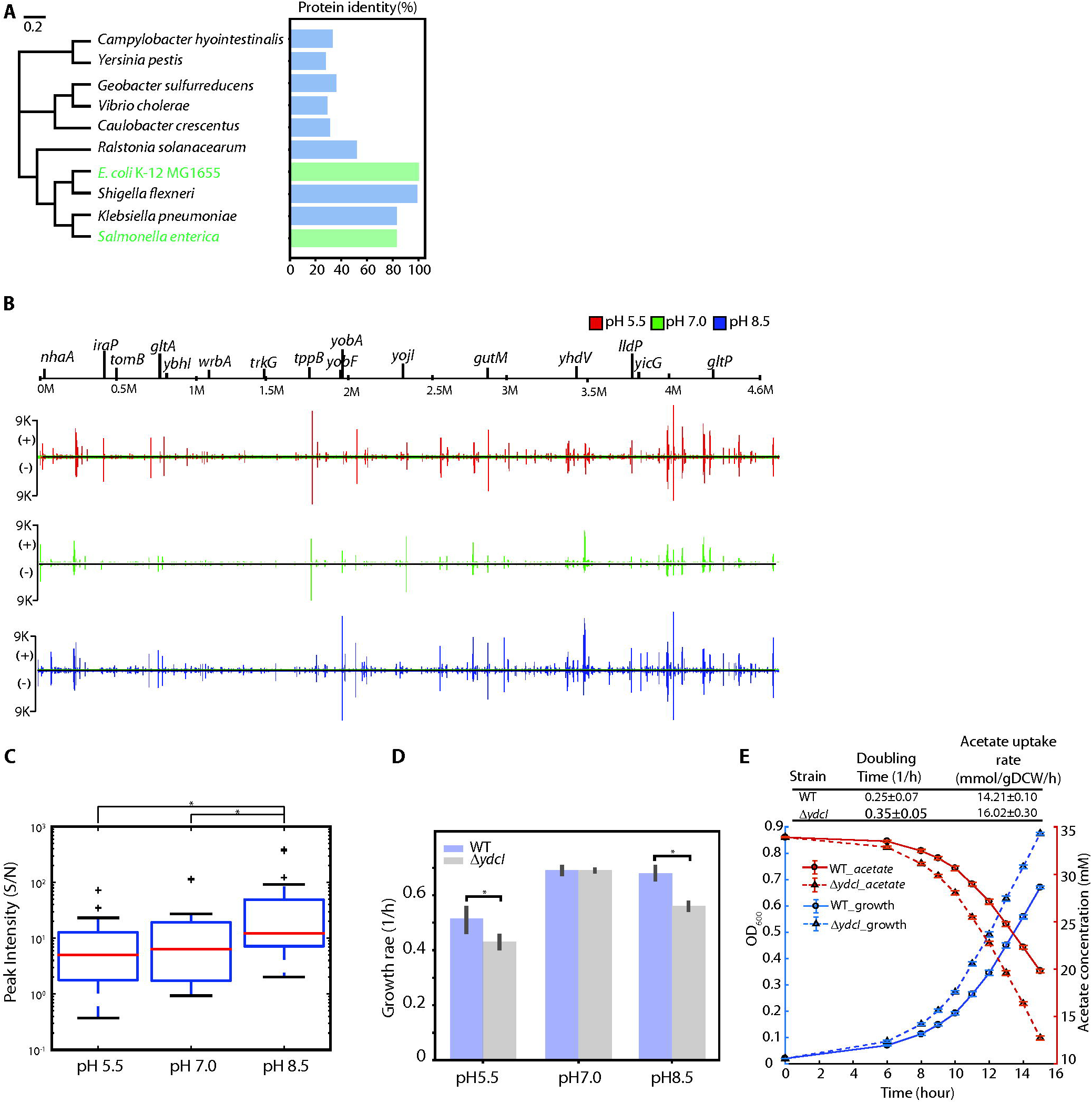
The regulatory role of the uncharacterized TF YdcI is involved in proton and acetate metabolism in *E. coli* K-12 MG1655. (A) Phylogenetic trees displaying the relatedness of YdcI from *E. coli* K-12 MG1655 and from *Salmonella enterica*. (B) Genome-wide YdcI DNA binding. YdcI binding across the genome was compared under different pH conditions in the *E. coli* K-12 MG1655 by ChIP-exo. (C) Peak intensity (Signal/Noise) of YdcI ChIP-exo binding sites at pH 5.5, pH 7.0, and pH 8.5. Among three different pH conditions, peak intensity was strongest at pH 8.5 (* indicates ranksum test *p*-value <0.05). (D) Growth rate of wild type and *ydcI* deletion strain at low pH, neutral pH, and high pH media. (E) Growth and acetate uptake rates of wild type and *ydcI* deletion strains in acetate growth medium. Measurement of acetate uptake rate of wild type and *ydcI* deletion strain.

To test our hypothesis, we conducted ChIP-exo experiments for YdcI at different pH conditions (Figure 5B). Under low pH conditions, YdcI bound to 16 locations, and two-thirds of these binding peaks were found in intergenic regions. Under neutral or high pH conditions, YdcI bound to all sites identified at low pH conditions but had differential binding intensity. Thus, we further analyzed the ratio of signal to noise (S/N) and found that YdcI had the highest average binding intensity at high pH medium (Figure 5C). More important, we found that four of the target genes, *nhaA, dtpA, lldP,* and *gltP*, encode proton transporters, which play important roles in the acid stress response. Therefore, we decided to examine the growth phenotype of the *ydcI* deletion strain at low pH, neutral pH, and high pH media (Figure 5D). At pH 5.5, the *ydcI* deletion strain showed significant growth defects compared to the wild type. However, there was no defect observed at neutral pH conditions. These data confirmed that YdcI is essential to maintain physiological activity at low pH conditions and to regulate the proton transfer in *E. coli*.

YdcI has another important binding site at the gene *gltA*, which encodes a citrate synthase in *E. coli* K-12 MG1655. It is induced and becomes the rate-limiting step for the TCA cycle when acetate is the sole carbon source (68, 69). A previous study hypothesized that YdcI may regulate the carbon flux in the TCA cycle through *gltA* expression (70). To test this hypothesis, we compared the growth of *E. coli* WT and the *ydcI* deletion strain in acetate medium (Figure 5E). The *ydcI* deletion strain grew significantly faster than the wild type, showing that YdcI represses the gene *gltA*. We verified that the acetate uptake rate increased upon *ydcI* deletion compared to WT using high performance liquid chromatography (HPLC), which confirmed that YdcI is also involved in regulating the carbon flux in the TCA cycle.

### Case III: YeiE is a transcription factor that iron homeostasis

Group III includes candidates with neither biochemical characterization nor biological function prediction. Among them, the global binding profile of LysR-type YeiE showed over 100 binding sites across the *E. coli* K-12 MG1655 genome (Supplementary Figure S9) (6). Target genes of YeiE are involved in diverse biological processes, including transport and metabolism, cell wall / membrane biogenesis, signal transduction, and transcriptional regulation (Figure 6A). In addition, functional classification showed that approximate 42% (43/102) of YeiE bindings is involved in main transport processes, including amino acids, carbohydrate and inorganic ions, though it is not significantly enriched in any functional group. This data suggests that YeiE may play multiple biological roles in *E. coli* K-12 MG1655. To further investigate the potential functions of YeiE, we examined the expression profiles of the ΔA*yeiE* strain. Three COG functional groups were significantly (*p*-value < 0.01) associated with the functions of YeiE: energy production and conversion, amino acid transport and metabolism, and inorganic ion transport and metabolism (Supplementary Figure S10). Notably, we found that many metal ion homeostasis-related genes, such as *entS*, *entC*, *cirA*, *fhuA*, *fhuF*, *fepB,* and *feoA*, were down-regulated in the *yeiE* deletion strain (Figure 6B). These results suggest that YeiE may be involved in the iron-uptake regulation pathway.

**Figure 6.**
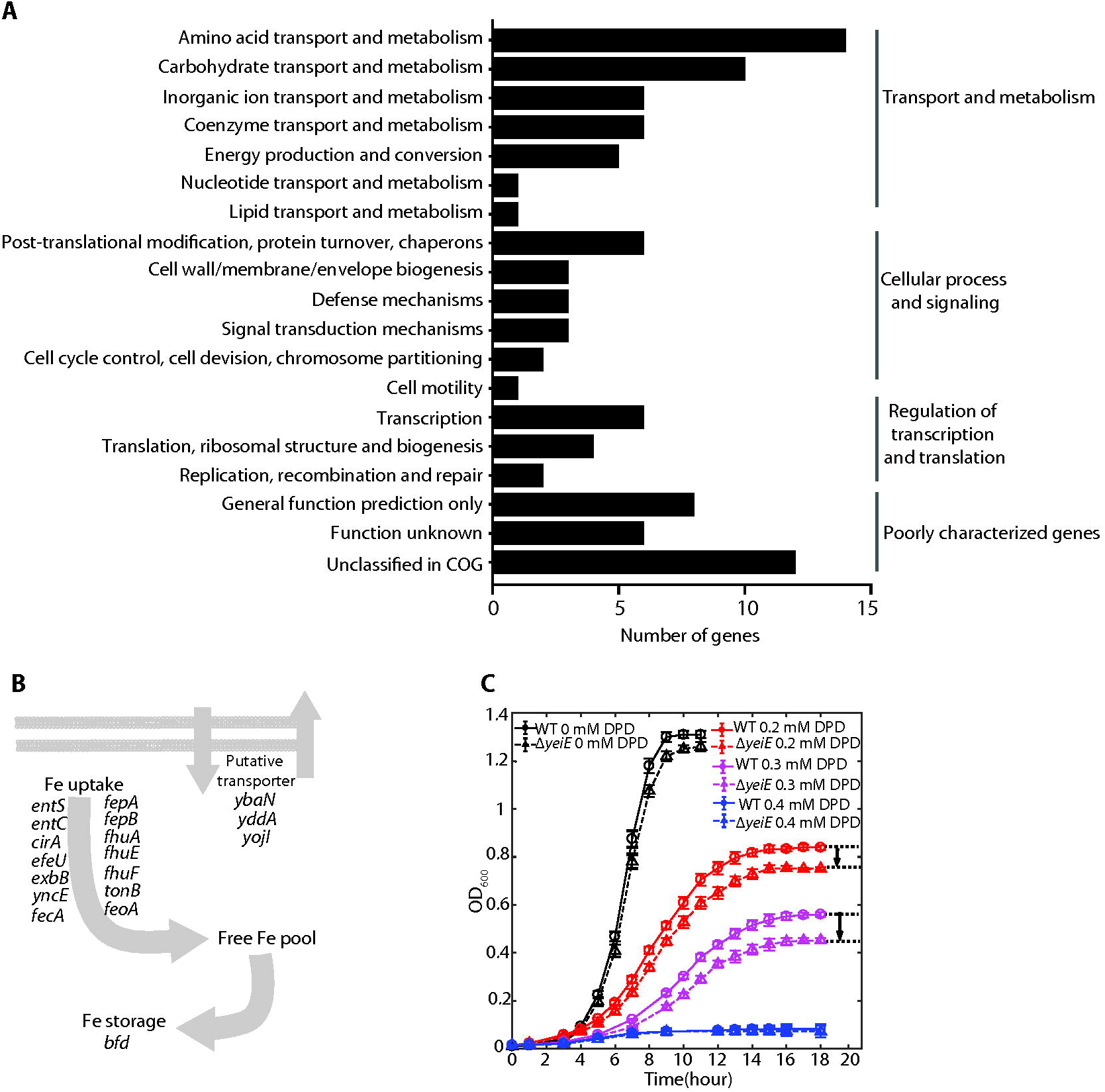
The regulatory role of the uncharacterized TF YeiE is involved in maintaining iron homeostasis under iron limited conditions in *E. coli* K-12 MG1655. (A) Functional classification of target genes from YeiE genome-wide bindings. The enriched functions are in three groups: amino acid transport / metabolism, carbohydrate transport / metabolism, and inorganic ion transport / metabolism. (B) The proposed regulation roles of genes down-regulated in the *yeiE* deletion strain. (C) Growth profile of wild-type and *yeiE* deletion strain in iron-free M9 minimal medium supplemented with 0, 0.2 mM, 0.3 mM, 0.4 mM 2,2’-dipyridyl (DPD), an iron chelator, respectively.

To examine the role of YeiE in inorganic ion transport and metabolism, we measured the growth profiles of the wild type and *yeiE* deletion strain in M9 medium with or without iron chelator (Figure 6C). There was no appreciable difference between the growth profiles of the two strains in the iron-rich condition without iron chelator. Then we added iron chelator (2,2’-dipyridyl, DPD) to the final concentration 0.2 mM. When an iron chelator (0.2 mM 2,2’-dipyridyl, DPD) was introduced, the *yeiE* deletion strain grew slower than the wild type in the early-mid log phase. As cells entered into late log phase, different growth rates were observed. The differences between the wild-type and *yeiE* deletion strain increased with the concentration of iron chelator in the media. When the concentration of iron chelator reached 0.4 mM, neither strain could enter the log phase. The fact that this growth defect was only observed under iron-limited conditions suggested that YeiE is involved in iron-uptake pathways under iron-limited conditions.

## DISCUSSION

The characterization of a transcriptional regulatory network (TRN) is an important step in our understanding of organism function and evolution. A major limitation toward this goal is that we do not have a complete set of characterized TFs for an individual organism as of yet. Here, we addressed this gap with the development of an integrated bioinformatic and experimental workflow. We applied this workflow to *E. coli* K-12 MG1655, one of the most well-studied organisms, and discovered ten previously uncharacterized TFs *in vivo*. We reconstructed the regulon for each of novel TFs and determined the physiological roles for three of them; YiaJ is involved in the utilization of L-ascorbate (Figure 7A), YdcI is involved in proton and acetate metabolism (Figure 7B and 7C), and YeiE is involved in iron uptake under iron-limited conditions (Figure 7D). For other candidates with binding sites, it is worth doing further exploration about the regulatory roles. In addition, we found that *in vivo* binding patterns of YbiH and YbaO were consistent with the genomic SELEX results, though genome wide binding profile of YbiH showed some extra target genes (Supplementary Figure S7) (71, 72). That suggested that the binding patterns of some regulators are very consistent between *in vivo* and *in vitro* methods. We note that while ChIP-exo is commonly used for the mapping of TF-DNA interactions, its application to the elucidation of regulon function is limited by the knowledge of suitable conditions that activate a target TF. Our results have several important implications.

**Figure 7.**
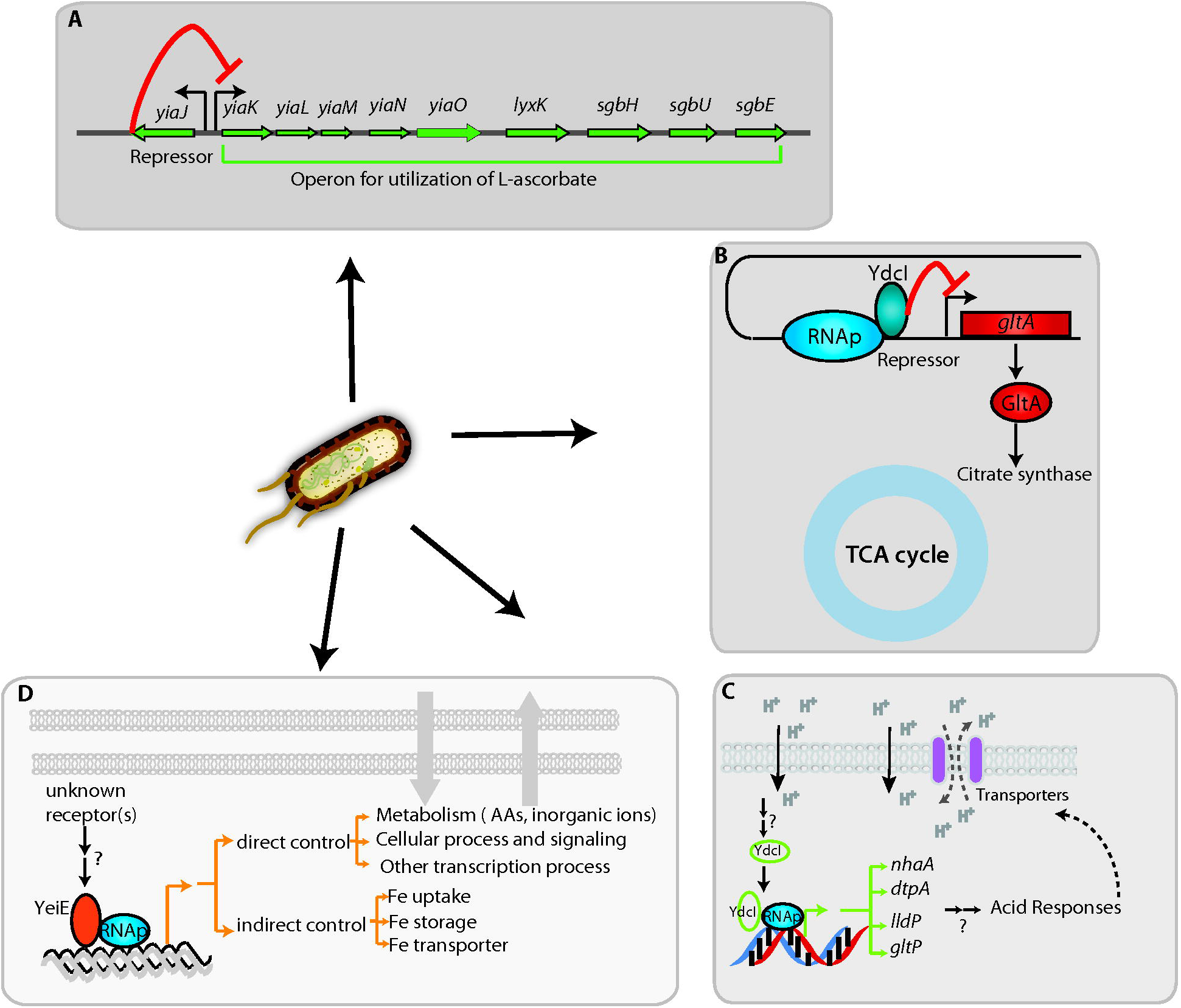
The model for the regulatory network integrating three candidate TFs (YiaJ, YdcI, and YeiE) and their biological functions in *E. coli* K-12 MG1655. (A) YiaJ is a regulator that controls the operon *yiaK*-*yiaL*-*yiaM*-*yiaO*-*lyxK*-*sgbH*-*sgbU*-*sgbE* in the catabolism pathway. (B) YdcI inhibits the transcription of target gene *gltA*, resulting in down-regulation of citrate synthase that is required in the TCA cycle. (C) YdcI binds to genomic DNA and activates target genes *nhaA*, *dtpA*, *lldP*, *gltP* that are responsible for proton transfer. (D) YeiE affects multiple transporters (amino acids, inorganic ions, lipids) and metabolic processes, maintaining iron homeostasis at iron limited conditions.

First, the ten newly identified TFs represent a 6% increase to the 185 already known TFs. Furthermore, we provided new knowledge about the co-binding of candidate TFs and known TFs (Supplementary Figure S6). This TF discovery workflow enables the systematic examination of the remaining putative TFs identified by the initial computational step of the workflow. In this study, six of the examined candidate TFs were not found to have any binding sites at test conditions. This failure to identify binding sites could have happened for two reasons: 1) our conditions did not activate these TFs; e.g., YagI was recently identified as a regulator of xylonate catabolism using the SELEX method *in vitro* (73); and, 2) current prediction algorithm methods may generate false positive candidates. Recently, we found that YihY and YchA were annotated as putative inner membrane protein and transglutaminase-like / TPR repeat-containing protein, respectively, though their physiological functions remain unclear. However, it is still necessary to develop a systematic workflow to predict and validate TFs and improve our knowledge of the TRN.

Second, differential expression data between wild type and uncharacterized TF deletion strains allowed us to reconstruct new regulons. We reconstructed a regulatory network containing 47 new regulatory interactions between candidate TFs and their target genes (Supplementary Figure S6). Specifically, we added more regulatory information for 25 target genes that previously had no known regulator. The reconstructed regulons suggest functional associations between both characterized and uncharacterized genes (Supplementary Figure S11). For instance, as a periplasmic protein, the physiological role of GltF in *E. coli* is still unknown. Functional enrichment suggests that it may transport inorganic ions or other metabolites. Future experimental studies are needed to discover the functions of these uncharacterized genes.

Third, detailing the functions of three of the ten regulons adds to our understanding of the TRN in *E. coli*. Iron response is a key characteristic in most enterobacteria, as well as bacteria in general. Although Fur is a well known TF for iron response, the discovery of YeiE as an active TF under low iron conditions adds to our understanding of the overall iron response (Figure 7D). Low iron levels are especially important in understanding the interactions between pathogens and hosts (74, 75). Transcriptional regulation of ascorbate metabolism has been largely unknown, and the discovery of the role that YiaJ plays helps fill this knowledge gap (Figure 7A). The transcriptional repressor YiaJ belongs to the IclR family and controls the hypothetical ascorbate transport system (named *yiaMNO*) and four genes (*lyxK-sgbH-sgbU-sgbE*) encoding ascorbate catalytic enzymes (6, 76).

In this study, we have presented a workflow for the systematic discovery of uncharacterized TFs, thus enabling the reconstruction of their regulons. A study of an initial set of 16 candidate TFs demonstrated that the workflow can systematically elucidate TF functions in *E. coli*. This workflow also provides a path for discovery of uncharacterized gene functions that were found in the newly discovered regulons. Given that many TFs are broadly conserved in gram-negative strains, this workflow can be applied to other bacteria, thus allowing a study of the similarities and differences in gene regulation between species (77). Such characterization would offer an unprecedented insight into the function and evolution of regulatory networks in bacteria.

## DATA AVAILABILITY

The whole dataset of ChIP-exo and RNA-seq has been deposited to GEO with the accession number of GSE111095.

## AUTHOR CONTRIBUTIONS

Y.G., D.K. and B.O.P. designed the study. Y.G.and D.K. performed experiments. J.T.Y., A.D. and J.E. developed the computational algorithm and performed analysis. Y.G., D.K.,S.W.S.,I.K., A.V.S. and X.F. did data analysis. K.C. and N.M. contributed to the protein structure analysis. Y.G., J.T.Y., B.K.C., D.K. and B.O.P. wrote the manuscript, with contributions from all other authors.

## ACKNOWLEDGEMENT

We thank Richard Szubin for help with ChIP-exo and RNA-seq library sequencing. We thank Zachary King, Justin Tan and Amitesh Anand for helpful discussions. We thank Marc Abrams for reviewing and editing the manuscript.

---

**Table 1. Category of uncharacterized transcription factors in this study**

